# HiFi Metagenomic Sequencing Enables Assembly of Accurate and Complete Genomes from Human Gut Microbiota

**DOI:** 10.1101/2022.02.09.479829

**Authors:** Chan Yeong Kim, Junyeong Ma, Insuk Lee

## Abstract

Advances in metagenomic assembly have led to the discovery of genomes belonging to unculturable microorganisms. Metagenome-assembled genomes (MAGs) often suffer from discontinuity and chimerism. Recently, nanopore metagenomic sequencing assembled 20 complete MAGs (cMAGs) from 13 human fecal samples, but with low nucleotide accuracy. Here, we report 102 cMAGs obtained by high-accuracy long-read (HiFi) metagenomic sequencing of five human fecal samples, whose initial circular contigs were filtered for authentic prokaryotic genomes using our bioinformatics workflow. Nucleotide accuracy of the final cMAGs was similar to that of Illumina sequencing. The cMAGs could exceed 6 Mbp and included complete genomes of diverse taxa, including entirely unculturable RF39 and TANB77 orders, whose genomes have not been characterized yet. Moreover, cMAGs revealed that regions hard to assemble by short-read sequencing comprised mostly genomic islands and rRNAs. HiFi metagenomic sequencing will facilitate cataloging accurate and complete genomes of human gut microbiota, including unculturable species.

## Introduction

Despite advances in culturomics technology, most human gut prokaryotic species remain unculturable^1-3^. Therefore, conventional cataloging of microbial genomes, based on isolation of clonal genomic DNA followed by sequencing and assembly, may not be applicable to all human gut commensals. *De novo* metagenomic assembly using human fecal sequencing samples has proven useful in reconstructing the genomes of gut species including unculturable taxa^2,3^. Nevertheless, these metagenome-assembled genomes (MAGs) are generally discontinuous and chimeric because of conserved, repetitive, and mobile sequences^4,5^. Long-read metagenomic sequencing using Oxford nanopore technology (ONT) has enabled the assembly of 20 circularized complete MAGs (cMAGs) from 13 human stool samples, albeit with low nucleotide accuracy^6^. More recently, PacBio high-accuracy long-read (HiFi) sequencing, which has become popular for the assembly of reference animal and plant genomes^7,8^, has been applied for the analysis of complex microbiomes, such as sheep fecal samples^9^ and chicken cecum samples^10^. HiFi repetitive sequencing of a circularized SMRTbell library calls reads by consensus, substantially improving nucleotide accuracy while maintaining long read length. Moreover, specialized assemblers for HiFi metagenomic assembly^10,11^ enable the highly accurate reconstruction of cMAGs.

In the present study, we conducted an exhaustive assembly of HiFi metagenomic sequencing reads from five human fecal samples. Seeking only circularized contigs, we skipped the binning procedure. We developed a bioinformatics workflow to filter initially assembled circular contigs for authentic complete prokaryotic genomes. Eventually, we obtained 102 cMAGs representing the complete genomes of diverse phylogenetic groups of human gut microbiota. We demonstrate that cMAGs obtained by HiFi metagenomic sequencing share similar nucleotide accuracy as Illumina short-read sequences. Several cMAGs exceeded 6 Mbp, with the longest reaching 6.77 Mbp, larger than 99% of all human gut prokaryotic species in terms of genome size. We also obtained cMAGs for the unculturable RF39 and TANB77 orders, for which no genomes have yet been isolated and reported. Alignment of genomes to HiFi cMAGs revealed that regions hard to assemble by short-read sequencing comprised mostly genomic islands and rRNAs. These results suggest that HiFi metagenomic sequencing can be used to assemble accurate and complete genomes of human gut microbiota, including unculturable species.

## Results

### HiFi metagenomic sequencing assembles cMAGs belonging to diverse gut microbiota taxa

We obtained public HiFi metagenomic sequencing data from four human fecal samples^12^. Two samples were pooled from vegan donors and the other two from omnivore donors. In addition, we produced in-house HiFi metagenomic sequencing data based on a fecal sample from a healthy Korean omnivore donor using the Sequel II platform. Compared to a recently published ONT metagenomic sequencing dataset on human fecal samples^6^, the HiFi metagenomic sequencing data used in this study displayed similar total base pairs but much longer reads and significantly higher base quality **(Extended Data Fig. 1, Supplementary Table S1)**.

To obtain as many circular contigs as possible, we assembled HiFi sequencing reads for each sample using three different metagenomic assemblers, metaFlye^13^, HiCanu^11^, and hifiasm_meta^10^, which yielded 2,283, 481, and 590 circular contigs, respectively, or 3,354 in total. Because these circular contigs might contain viral genomes, plasmids, and misassembled closed contigs, we developed a bioinformatics workflow to select only authentic prokaryotic genomes **(Fig. 1a)**. First, we filtered circular contigs using biological priors and structure parameters, including sequence length ≥ 100 kbp, ≥ 100 genome taxonomy database (GTDB) marker proteins, presence of rRNAs, ≥ 20 tRNA types, and no assembly bubble or repeat **(Extended Data Fig. 2a, b)**. Although metaFlye initially assembled the largest number of circular contigs, ~97.7% (2,231) were filtered out using these criteria. Accordingly, 145, 76, and 52 circular contigs passed the first filtering step by hifiasm_meta, HiCanu, and metaFlye, respectively **(Extended Data Fig. 2c)**. Because we applied multiple assemblers on pooled fecal samples, any resulting redundant contigs with near-identical sequences (average nucleotide identity, ANI > 0.99 and maximum alignment coverage > 0.95) were removed after calculating all pairwise genome similarities **(Extended Data Fig. 2d–g, Supplementary Table S2)**.

**Fig. 1.**
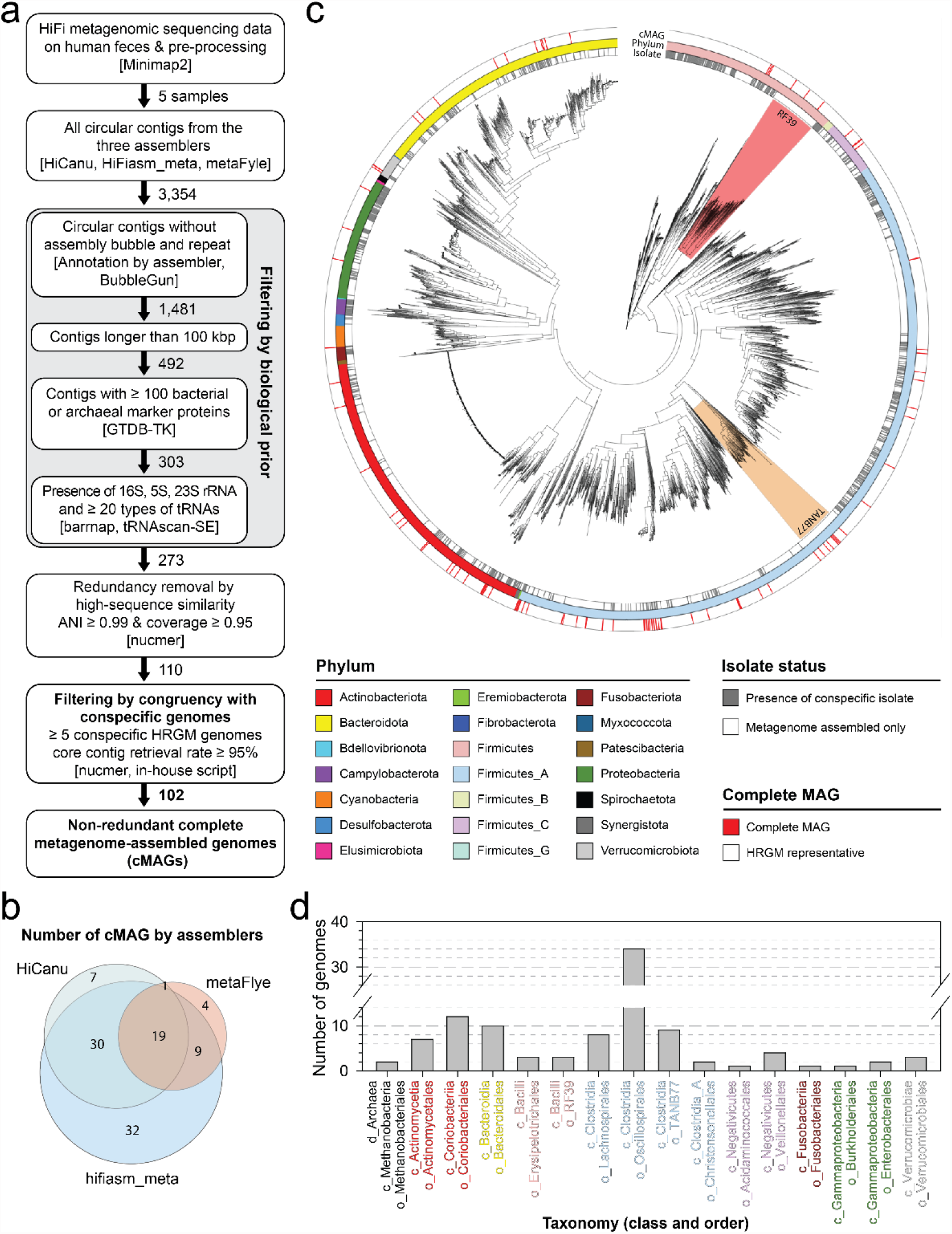
Assembly of 102 complete metagenome-assembled genomes (cMAGs) from human fecal high-accuracy long-read (HiFi) metagenomic sequencing samples. **a**, Schematic diagram illustrating the HiFi cMAG assembly workflow. **b**, Number of final cMAGs assembled from each assembler. Overlapping areas represent redundant genomes assembled by more than one assembler. **c**, Maximum likelihood phylogenetic tree of 5,486 bacteria from the human reference gut microbiome (HRGM) and HiFi-assembled cMAGs. RF39 and TANB77 clades are highlighted in red and orange, respectively. Color bars from the inner to the outer circles represent isolated status, phylum, and HiFi cMAGs, respectively. **d**, Number of cMAGs for each phylogenetic order. The text color represents phylum classification for each order. The plot uses the same color codes as in (c) except for the archaeal order Methanobecteriales (black).

Repeated sequences could cause early assembly closing, generating prokaryotic genomes with significant gaps^14^. These defective genomes may not be detected even by alignment with conspecific genomes, unless isolated genomes are available for the species. Therefore, we designed another filtering step based on the congruency of each circular contig with its conspecific genomes from the human reference gut microbiome (HRGM)^3^. We hypothesized that although individual HRGM genomes might not be complete, their core contigs shared among most conspecific genomes likely originated from genuine species. Therefore, for each circular contig, we first identified conspecific HRGM genomes and determined their core contigs. Then, we aligned the core contigs to the query circular contig, and excluded the latter if the retrieval rate of core contigs was < 95% or there were fewer than five conspecific genomes **(Extended Data Fig. 3a, b, Supplementary Table S3)**. Circular contigs that did not meet these criteria were significantly shorter than their conspecific isolated genomes **(Extended Data Fig. 3c)** and contained significant gaps compared to their closest isolated genome **(Extended Data Fig. 3d–f)**. Overall, our bioinformatics workflow retained 102 of the 3,354 circular contigs initially assembled.

Previously, a skewed GC content was proposed as a metric for verifying the correctness of assembled prokaryotic genomes^15^. We manually examined the cumulative GC-skew pattern of the 102 circular contigs and classified them into five classes: very clear, clear, decent, poor, and no pattern **(Supplementary Table S4)**. Unexpectedly, many circular contigs with poor or no GC symmetry patterns possessed the characteristics of genomes from specific lineages rather than misassembled ones. For example, five out of six Oscillospiraceae family genomes showed poor or no GC-skew pattern, and all *Gemmiger* genus, *Bifidobacterium adolescentis*, and *Bifidobacterium longum* genomes showed no GC symmetry **(Extended Data Fig. 4)**. Indeed, the latter two were previously reported as not having GC symmetry^15^. Accordingly, we decided not to filter out circular contigs based on the GC-skew pattern.

The final 102 cMAGs could represent the complete genomes of human gut prokaryotic species. Notably, hifiasm_meta contributed most of the 102 cMAGs (~88.2%) **(Fig. 1b)**. Although bias toward a few bacterial clades (e.g., Oscillospirales) was observed, cMAGs from HiFi metagenome sequencing reads covered diverse phylogenetic groups of human gut microbiota. Based on the GTDB annotation, we obtained cMAGs for both Archaea and Bacteria, comprising 9 phyla, 11 classes, 14 orders, 24 families, 52 genera, and 84 species **(Fig. 1c, d, Supplementary Table S5)**.

### The cMAGs from HiFi metagenomic sequencing reads are accurate and can be large

Next, we evaluated the nucleotide accuracy of cMAGs obtained by HiFi metagenomic sequencing. Even without taking into account sequencing errors, same-strain prokaryotic genomes assembled from different samples are hardly identical to each other because of genetic variation accrued via horizontal gene transfer and spontaneous mutation. To assess the nucleotide accuracy of cMAGs, we chose *Bifidobacterium animalis*, which showed significantly lower genetic variation than other gut commensal species. *B. animalis* (HRGM_Genome_1769) displayed the tenth lowest single nucleotide variation (SNV) rate among the 1,521 HRGM species with known SNV values and the lowest SNV rate among HRGM species with isolated genomes^3^ **(Fig. 2a)**. We then aligned the *B. animalis* HiFi cMAG (OMN01_MFL_0491) to its closest isolated genome in RefSeq^16^ (GCF_000224965.2). Interestingly, the two genomes were nearly identical (ANI > 0.9999 and alignment coverage > 0.9999) **(Fig. 2b)**, even though they were assembled at different locations and time points (USA in 2021 and Italy in 2013, respectively). When we compared the SNV density of *B. animalis* HiFi cMAG to that of other *B. animalis* genomes in HRGM **(Extended Data Fig. 5)**, we did not observe a substantial deviation in the SNV rate (maximum 14.6%) **(Fig. 2c)**, suggesting that SNVs were attributable mostly to spontaneous mutations and errors occurring during short-read sequencing. To generalize these findings, we explored the percentile rank of the SNV rate among 77 HiFi cMAGs in their conspecific genomes. The 50th percentile rank fell within the interquartile range of the boxplot, and only four cMAGs were above the 99th percentile rank **(Fig. 2d)**. Hence, SNV rates of most HiFi cMAGs do not deviate from the distribution of their conspecific HRGM genomes.

**Fig. 2.**
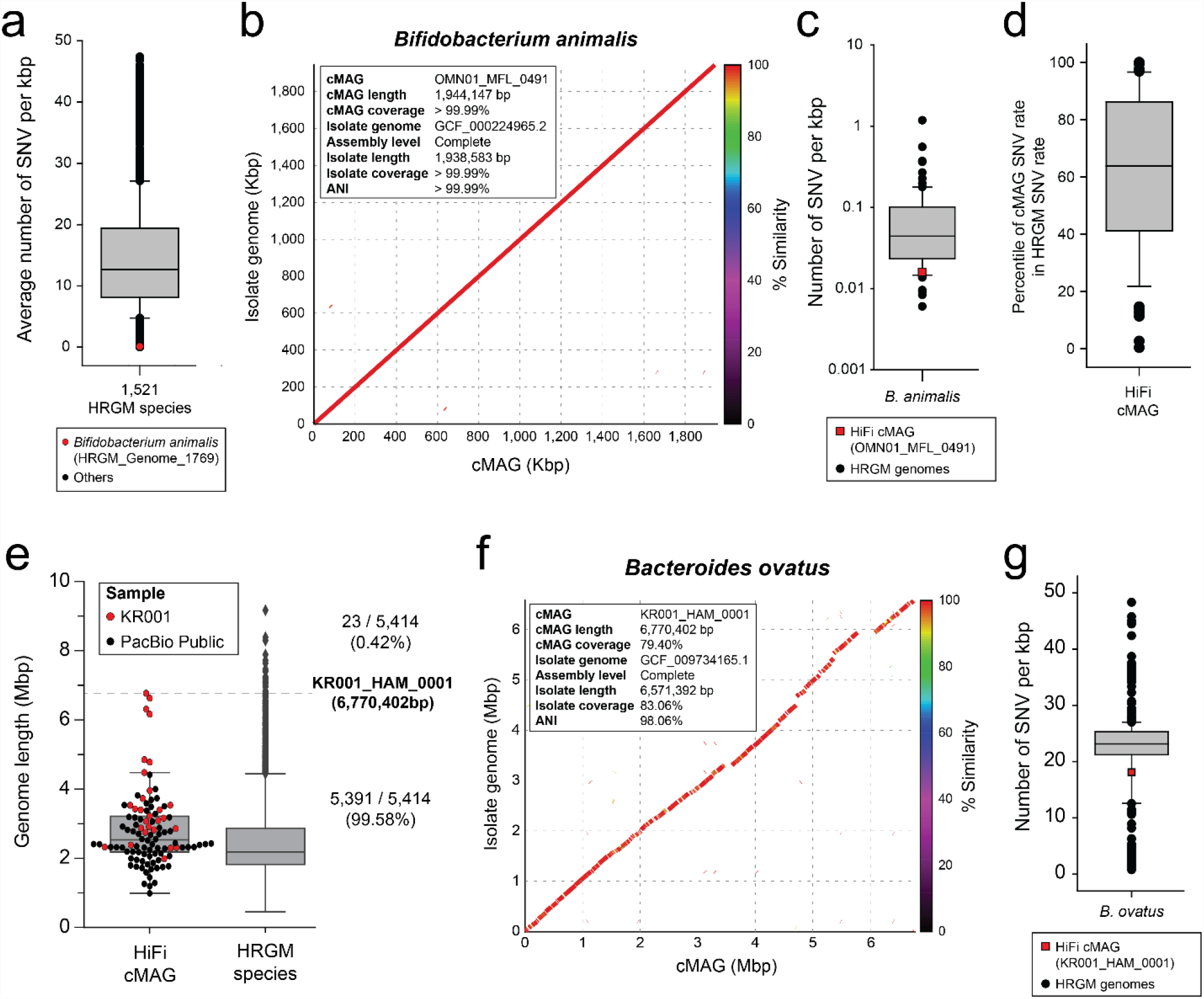
HiFi metagenomic sequencing reconstructs accurate and long prokaryotic genomes. **a**, Single nucleotide variation (SNV) rate of 1,521 HRGM species and *Bifidobacterium animalis* (red dot). **b**, Genome alignment plot between *B. animalis* cMAG (x-axis) and the closest isolated genome (y-axis). **c**, SNV rate of *B. animalis* cMAG and its conspecific HRGM genomes. **d**, Percentile rank SNV rate of cMAGs among conspecific genomes. **e**, Assembled genome length of cMAG and HRGM species. The horizontal dashed line represents the length of the longest HiFi cMAG. **f**, Genome alignment plot between *Bacteroides ovatus* cMAG (x-axis) and the closest isolated genome (y-axis). **g**, SNV rate of *B. ovatus* cMAG and conspecific HRGM genomes.

We next examined the size achieved by HiFi metagenomic sequencing. Four cMAGs in the *Bacteroides* genus were over 6 Mbp and the longest seven cMAGs were assembled from the Korean sample (KR001), which had the greatest sequencing depth **(Fig. 2e, Supplementary Table S1)**. The longest cMAG (6,770,402 bp) was KR001_HAM_0001, which might correspond to *Bacteroides ovatus* and is larger than 99.58% of 5,414 HRGM species in terms of genome size. The longest cMAG previously reported by ONT metagenomic sequencing was 3,825,229 bp^6^. Thus, KR001_HAM_0001 is the largest cMAG ever published. To confirm the integrity of KR001_HAM_0001, we compared it with the closest isolated genome (RefSeq: GCF_009734165.1), to which it was highly similar in most genomic regions (ANI > 0.98 and alignment coverage ~0.80) **(Fig. 2f)**. In addition, the SNV rate of KR001_HAM_0001 did not diverge significantly from that of its conspecific HRGM genomes (14% from the lowest) **(Fig. 2g)**. These results suggest that HiFi metagenomic sequencing assembles accurate and complete genomes of human gut microbiota, including species with a genome size exceeding 6 Mbp.

### HiFi metagenomic sequencing assembles complete genomes for unculturable taxa

More than 80% of human gut species are thought to be unculturable^2,3^. Anticipating the benefit of genome cataloging through cMAGs, we surveyed previously entirely unculturable human microbial taxa. The culturability status of each cMAG was defined based on the presence of isolated conspecific genomes in the HRGM, hGMB, and NCBI genome databases. This yielded 63 culturable cMAGs and 39 unculturable cMAGs **(Fig. 3a, Supplementary Table S4)**. Among the 63 culturable cMAGs, 24 had discontinuous genomes with gaps and consisted of genomic scaffolds. Therefore, cMAGs obtained by HiFi metagenomic sequencing improved genome quality. Applying GTDB annotations, we identified cMAGs for unculturable taxa comprising 35 species, 19 genera, 4 families, and 2 orders **(Fig. 3b, Supplementary Table S5)**.

**Fig. 3.**
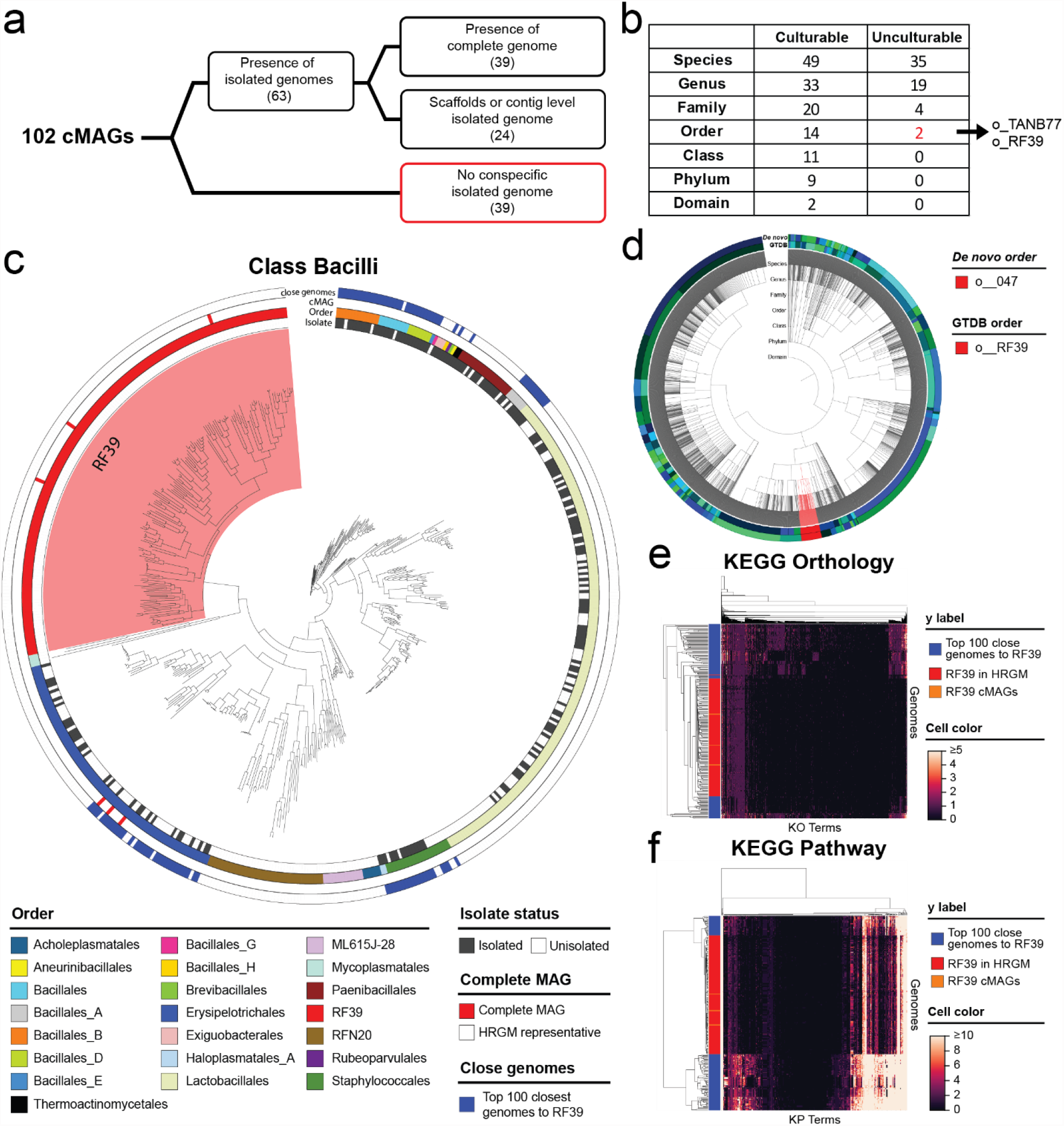
Assembly of complete genomes for unculturable microorganisms. **a**, Number of cMAGs with isolated and complete conspecific genomes. The 39 cMAGs for unculturable taxa are marked in red. **b**, Number of culturable and unculturable taxa among the 102 cMAGs. **c**, Maximum likelihood phylogenetic tree of HRGM species and HiFi cMAGs in the Bacilli class. The red-highlighted area represents the RF39 order. Isolated status, order, HiFi cMAG, and top 100 closest genomes to the RF39 order are annotated from the innermost to the outermost circle, respectively. **d**, Phylogenetic tree of 4,545 bacterial species of HGM. The inner circle annotates the order according to the GTDB; whereas the outer circle represents the order according to the HGM *de-novo* classification. RF39 and o_047 orders are highlighted in red. **e, f**, Hierarchically clustered heatmaps representing KEGG Orthology (e) and KEGG Pathway (f) profiles of 157 RF39 (red and orange rows) and the top 100 closest genomes to RF39 (blue rows).

Although the RF39 and TANB77 orders include 154 and 120 species, respectively (based on the HRGM), none of the species belonging to these taxa have been isolated so far. We identified three cMAGs belonging to RF39 and nine to TANB77. HiFi metagenomic sequencing assembled complete genomes for these large and entirely unculturable taxa from the human gut for the first time. RF39 is a newly defined order of the Bacilli class, with more than 50% of species being classified neither at order level nor through consensus with NCBI taxonomic annotation **(Extended Data Fig. 6a)**. Here, RF39 formed a distinct clade in the phylogenetic trees of Bacteria and Bacilli **(Fig. 1b and Fig. 3c)**. Given that the phylogenetic tree and GTDB classification were generated based on the same bacterial marker proteins (bac120), we reviewed the *de novo* classification using an independent set of marker proteins^17^. Interestingly, all RF39 bacteria matched the *de novo* order o_047, and *vice versa* **(Fig. 3d)**. Next, we predicted the microbial proteins of species belonging to the RF39 order and their neighboring species, and performed hierarchical clustering based on protein functional annotation. Most species of the RF39 order clustered together, supporting their functional distinctiveness **(Fig. 3e, f)**. TANB77 is another newly defined GTDB order, whose species are often classified into Clostridiales order by NCBI taxonomy **(Extended Data Fig. 6b)**. Using the same procedure as for RF39, we confirmed the functional distinctiveness of the TANB77 order **(Extended Data Fig. 6c–f)**. Together, these results imply that HiFi metagenomic sequencing enables the reconstruction of complete genomes belonging to unculturable prokaryotic clades of human gut microbiota.

### HiFi cMAGs reveal that hard-to-assemble genomic regions are mostly genomic islands and rRNA

MAGs obtained by short-read metagenomic sequencing contain many gaps that represent hard-to-assemble genomic regions. Repetitive and mobile sequences negatively affect correct and continuous assembly. Given that cMAGs are circularized genomes with no gaps, alignment of conspecific genomes generated by short-read metagenomic sequencing allows the systematic search for hard-to-assemble regions. Owing to the taxonomic diversity of the 102 cMAGs identified in this study, an unbiased taxonomic search was conducted. Quantification of genome coverage by cMAGs using conspecific HRGM MAGs (i.e., their contigs) revealed uneven coverage and the presence of regions with particularly poor coverage **(Fig. 4a)**. With 20% intervals for bins of genome coverage, most 1-kbp regions belonged to either > 80% or ≤ 20% genome coverages **(Fig. 4b)**, indicating a clear separation between well and poorly covered regions.

**Fig. 4.**
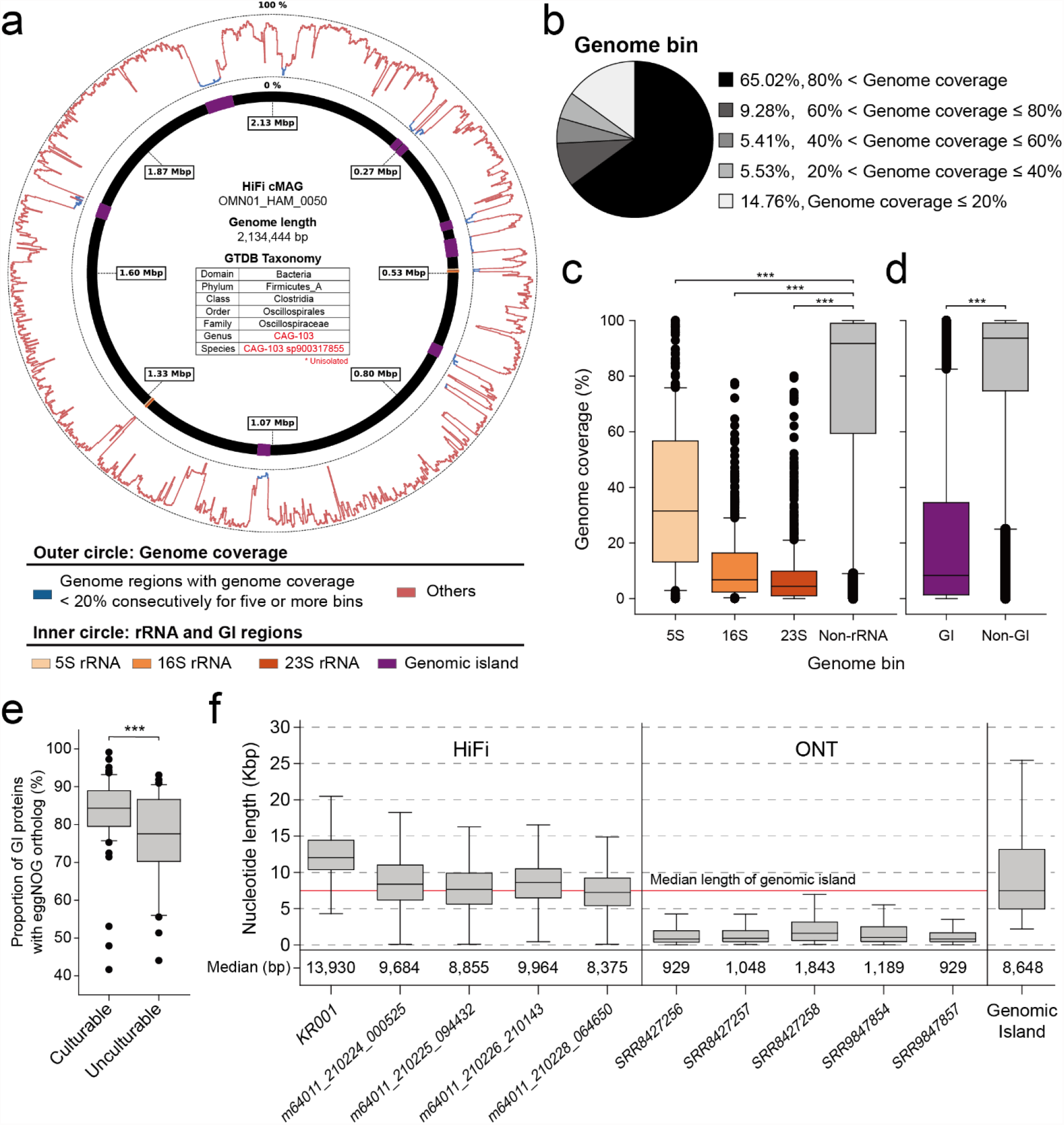
HiFi-assembled cMAGs retrieve hard-to-assemble regions by short-read assembly. **a**, Genome coverage and genome region annotation plot of a cMAG for the unculturable species OMN01_HAM_0050. The inner circle annotates rRNA and genomic island regions. The outer circle represents genome coverage by conspecific HRGM MAGs for every 1-kbp genome bin. **b**, Proportion of genome bins by genome coverage. **c, d**, Genome coverage of genome bins that include rRNAs (c) or genome islands (d). Genome coverage was compared by the two-sided Mann– Whitney U test. **e**, Proportion of genome island proteins with an eggNOG ortholog according to the culturability status of the host genome. The proportion was compared by the two-sided Mann–Whitney U test. **f**, Read length of long-read human fecal metagenomic samples and length of entire genome islands found among 102 cMAGs.

Highly conserved sequences (e.g., rRNAs) and mobile sequences (e.g., genomic islands) are notoriously difficult to retrieve from short-read metagenomic assembly^4,5^. Indeed, here, rRNAs and genomic islands aligned well with cMAG low-coverage regions (< 20% for more than five consecutive 1-kbp genome bins) **(Fig. 4a)**. Additionally, 1-kbp genome bins containing rRNAs and genome islands exhibited significantly lower genome coverage than other regions **(Fig 4c-d)**. Given that prokaryotic genomes have on average 4.2 copies of rRNAs^18^ and the relatively short length of these sequences, genomic island regions seem to be the leading cause of gaps in MAGs obtained by short-read sequencing. Genomic island proteins are more likely to be discovered from culturable taxa with isolated genomes. Thus, we hypothesized that cMAGs would provide opportunities to discover novel genome island genes. Indeed, genome island proteins of culturable taxa had a higher eggNOG ortholog^19^ annotation rate than unculturable taxa **(Fig. 4e)**. Moreover, the read length of HiFi metagenomic sequencing samples was similar or larger than the median length of genomic islands, whereas that of ONT metagenomic sequencing samples^6^ did not reach the median genome island length **(Fig. 4f)**. This suggests that HiFi reads likely cover genomic islands along with their adjacent regions, enabling correct mapping of foreign genetic elements to the host microbial genome.

## Discussion

HiFi sequencing offers substantial advantages in terms of base accuracy and read length^20^. In addition, HiFi metagenomic sequencing improves the quantity and quality of MAG assembly^9^. In the present study, we successfully retrieved complete circularized prokaryotic genomes of human gut microbiota through HiFi metagenomic sequencing without any binning process.

To take advantage of algorithmic complementarity, we utilized three different metagenomic assemblers: metaFlye, HiCanu, and hifiasm_meta. Initially, metaFlye generated three times more circular contigs than the other two assemblers, but most were filtered out, resulting in the lowest number of cMAGs. As metaFlye was not developed specifically for HiFi metagenomic sequencing, this may explain the high proportion of incomplete circular contigs. Instead, hifiasm_meta, which was developed to assemble HiFi metagenomic sequencing reads, alone retrieved > 88% of total cMAGs. Based on these results, we recommend hifiasm_meta for cataloging microbial genomes with HiFi metagenome sequencing.

Genome completeness is one of the most important criteria for cMAG. The completeness and contamination of MAGs are conventionally evaluated using single-copy genes^21^. This approach effectively filters out highly defective MAGs. However, single-copy gene-based completeness does not always coincide with actual completeness, especially for near-complete genomes. Therefore, we used the single-copy gene-based threshold to roughly filter out highly defective genomes, but not for the definitive evaluation of cMAGs. Moreover, lineage-specific marker proteins (e.g., checkM)^22^ sometimes underestimate MAG completeness, particularly for novel clades, due to the incorrect use of clade-specific marker genes, explaining why we used universal single-copy genes bac120 or arc122. Indeed, using Clostridiales lineage-specific markers for checkM assessment, we attained 80% cMAG completeness for TANB77 genomes. For the same reason, the genomes of many unculturable taxa showed relatively low completeness by checkM **(Supplementary Table S6)**. The ideal way to assess MAG quality is through comparison with reference-quality conspecific genomes. However, this is not always possible, particularly for unculturable taxa. Therefore, we devised a novel method based on congruency with conspecific MAGs, which effectively filtered out circular contigs with gaps.

No conspecific isolated genomes were found for 39 HiFi cMAGs because they belonged to unculturable taxa. Notably, even 24 HiFi cMAGs belonging to culturable taxa possessed conspecific isolated genomes consisting of discontinuous scaffolds. These results suggest that HiFi cMAGs can ameliorate existing human gut microbial genome catalogs not only for unculturable taxa but also for cultured ones. Many unculturable taxa have recently been defined through MAGs and have been newly classified by the GTDB^23^. These taxa often suffered from discordant annotation between the GTDB and NCBI classification systems. Indeed, the RF39 and TANB77 orders displayed distinct sets of marker proteins and functional profiles, supporting the GTDB as a reliable classification for unculturable taxa entirely composed of MAGs.

HiFi cMAGs enabled the unbiased examination of genomic regions poorly assembled by short-read assembly. The latter contained highly conserved sequences (e.g., rRNA) and mobile sequences (e.g., genome islands), which likely caused fragmentation during MAG assembly. Highly conserved regions of 16S rRNA complicate assembly, especially of the short-read type. Even MAGs with high completeness often lack 16S rRNA regions. Genomic islands are important because they confer strain diversity as their sequences originate from other species. Given that many binning algorithms cluster contigs based on species-specific k-mer frequency, genome islands rarely cluster together with other contigs originating from the same genome. Importantly, HiFi cMAGs can circumvent this problem because they do not rely on binning during assembly. In summary, we expect that HiFi metagenomic sequencing will facilitate not only cataloging accurate complete genomes of human gut microbiota but also the discovery of rRNA sequences and novel genes encoded in mobile genetic elements. Extensive application of HiFi metagenomic sequencing of human fecal samples would improve our understanding of the human gut microbiome. Although HiFi metagenomic sequencing is relatively expensive and requires abundant DNA (> 1.5 μg compared to ~0.1 μg for Illumina metagenomic sequencing), these limitations will be likely overcome by future technical improvements.

## Methods

### HiFi metagenomic sequencing data from human fecal samples

We collected four public HiFi metagenomic sequencing datasets of human fecal samples recently released by Pacific Bioscience^12^. Two of them were from pooled fecal samples of vegan donors, and the others were from pooled fecal samples of omnivore donors. In addition, we generated in-house HiFi metagenomic sequencing data from a fecal sample provided by a Korean donor following full informed consent and approval by the Yonsei University Institutional Review Board (IRB No. 4-2020-0309). The donor’s entire metagenomic DNA was sequenced by Macrogen Inc. (Republic of Korea). A high-quality and high-molecular weight genomic DNA sample, in which most fragments exceeded 40 kbp, was determined by pulsed-field gel or capillary electrophoresis and used for HiFi SMRTbell library preparation. The concentration of genomic DNA was measured by PicoGreen and its quality was evaluated in a Femto Pulse system (Agilent). We used 8 μg of input genomic DNA for HiFi library preparation. Femto Pulse helped determine the size distribution of genomic DNA < 15 kbp. Genomic DNA > 40 kbp was sheared by Megarupor3 and purified using AMPurePb magnetic beads. A HiFi SMRTbell library (10 μL) was prepared with the PacBio SMRTbell Express Template Prep Kit 2.0, annealed using the Sequel II Bind Kit 2.2 and Internal Control Kit 1.0.0, and sequenced with the Sequel II Sequencing Kit 2.0 and Sequel II SMRT cell 8M Tray. A 30-h video was recorded in each SMRT cell using the Sequel II platform. Subsequent procedures were performed according to the PacBio SampleNet-Shared protocol.

### Removal of host contaminants and *de novo* genome assembly of HiFi sequencing reads

We aligned all HiFi reads against the human reference genome (GRCh38.p13) with minimap2^24^ aligner using the -*ax asm20* parameter. Reads aligned to the human reference genome were considered human contaminants and were disregarded in downstream analysis. HiFi sequencing reads were assembled using three different metagenomic assemblers: HiCanu v2.1.1^11^, metaFlye v2.8.3-b1695^13^, and hifiasm_meta v0.2-r040^10^. For HiCanu, we used the -*pacbio-hifi* mode with recommended maxInputCoverage = 10000, corOutCoverage = 10000, corMhapSensitivity = high, and genomeSize = 3.7 M. For metaFlye, we applied the *--pacbio-hifi --meta* options. We ran hifiasm_meta with default parameters and used primary contigs.

### Filtering circular contigs based on biological priors

Only circular contigs obtained by metagenome-assembly entered our bioinformatics workflow to select cMAGs. We first filtered out ambiguous or non-prokaryotic circular contigs based on the assembly structure and prior biological knowledge. Contigs with assembly bubbles or repeats were excluded. The HiCanu assembler provides assembly bubbles and repeat annotations. For the contigs by metaFlye and hifiasm_meta assemblers, we identified repeats and bubbles from the output Graphical Fragment Assembly file using BubbleGun v1.1.1 software. Contigs shorter than 100 kbp were also removed, as they were highly unlikely to correspond to a complete prokaryotic genome **(Extended Data Fig. 2a)**. The remaining contigs were evaluated for completeness based on ubiquitous single-copy marker proteins. We predicted 120 bacterial (bac120) and 122 archaeal (arc122) marker proteins using GTDB-Tk v1.6.0^25^ and filtered out circular contigs with fewer than 100 of these marker proteins. Because marker counts gradually decreased before the threshold and dropped swiftly thereafter, faulty circular contigs could be distinguished **(Extended Data Fig. 2b)**. We also examined the existence of rRNAs (5S, 23S, and 16S rRNA) and tRNAs. We predicted rRNAs with barrnap v0.9 tools using the *--kingdom bac* and *--kingdom arc* parameters according to taxonomy by GTDB-Tk. We applied an evalue of 1e-04 for 5S rRNA (given its shorter length) and an evalue of 1e-06 (default) for the 16S and 23S rRNAs. We excluded contigs lacking rRNAs. We predicted tRNAs using tRNAscan-SE v2.0.7^26^ with *B* and *A* parameters for bacterial and archaeal contigs, respectively. We considered non-pseudo tRNAs only and disregarded circular contigs with fewer than 20 tRNA types. Circular contigs that passed each filtering step are listed in **Supplementary Table S3**.

### Removal of redundant circular contigs

The filtered circular contigs could be redundant because they were assembled from pooled samples. Therefore, we used sequence similarity to remove redundant circular contigs. We aligned every pair of circular contigs with nucmer 4.0.0beta2^27^ and used a delta-filter program to identify the best-bidirectional alignments. When we plotted the contig pairs by ANI (identical sequence length/aligned sequence length) and maximum alignment coverage (aligned sequence length/shorter contig length), we found a cluster of highly similar contigs with ANI > 0.99 and maximum alignment coverage > 0.95 **(Extended Data Fig. 2d)**. Most pairs within this group consisted of contigs from the same or the same type of sample **(Extended Data Fig. 2e)**. In addition, when we sorted the contig pairs by ANI multiplied by the maximum alignment coverage (similarity index), there was a clear discrimination between contig pairs above or below the similarity index of 0.9405 **(Extended Data Fig. 2f)**. Therefore, we determined ANI > 0.99 and maximum alignment coverage > 0.95 as the thresholds that removed redundant contigs while maintaining strain diversity. We selected a contig with the largest sum of ANI as a representative for each cluster of redundant contigs **(Extended Data Fig. 2g, Supplementary Table S2)**.

### Filtering circular contigs based on congruency with conspecific genomes

Filtered circular contigs by biological priors may still contain genomes with gaps. To identify such faulty genomes, we conducted the following processes using the HRGM catalog^3^: (i) finding conspecific HRGM genomes, (ii) identifying core contigs shared by most conspecific genomes, and (iii) aligning core contigs to the query circular contig **(Extended Data Fig. 3a)**. For each non-redundant circular contig, we found its conspecific genomes from the HRGM non-redundant genome set. We adopted a reduced search strategy because aligning all circular contigs against every non-redundant genome requires excessive computer power. We first aligned each circular contig against HRGM species representatives using nucmer with the *--mum* option followed by a delta-filter with *-r* and *-q* options. Only HRGM species representatives within the same genus as the circular contig by the GTDB taxonomy annotation were considered, because genomes of other genera were highly unlikely to meet the conspecific identity threshold (ANI > 0.95)^2,3,17,28,29^. Based on the species with the highest similarity index (ANI × maximum alignment coverage), we performed genome alignment using nucmer for all non-redundant HRGM genomes. By definition, HRGM conspecific genomes of the circular contig had ANI > 0.95 and maximum alignment coverage > 0.8. For three circular contigs (VEG02_HAM_0051, OMN01_HAM_0037, and OMN01_HAM_0001) within the *Collinsella* genus, which are reported to have an exceptionally high variant rate^3^, we adjusted the ANI threshold to 0.94 to find sufficient conspecific HRGM genomes for further analysis. We stopped searching when we found 100 conspecific genomes for each circular contig, while dismissing two circular contigs with fewer than five conspecific HRGM genomes.

Most conspecific genomes are MAGs, which are fragmented and may contain contamination. Here, we hypothesized that contigs shared by most conspecific MAGs (core contigs) were likely authentic genome sequences. Therefore, to find core contigs, we performed all pairwise genome alignments for the conspecific genomes of each circular contig using nucmer and delta-filter (same as above). For every contig of a conspecific genome, we considered that the contig was present in the other conspecific genome when more than half of the contig sequences were aligned with > 95% identity. A core contig was defined as longer than 5 kbp and present in more than 80% of conspecific genomes.

Finally, we aligned the core contigs to the query circular contig. Because the core contig set could contain redundant sequences, we used nucmer with the *--maxmatch* option. We calculated the core contig retrieval rate (aligned core contigs/core contigs) for each circular contig and excluded six circular contigs with core contig retrieval rate < 0.95 **(Extended Data Fig. 3b)**. A total of eight circular contigs (two with small conspecific genome count and six with low core contig retrieval rate) were excluded from the final list of cMAGs.

### Evaluation of cMAGs by the GC-skew pattern

For the final 102 cMAGs, we calculated and plotted the GC-skew pattern of each contig using the *gc_skew*.*py* script in the iRep^30^ package (1 kbp window, 10 bp slide). Next, we manually divided the cumulative GC-skew patterns into five classes: i) very clear (17 cMAGs), whereby the curve showed apparent symmetry with respect to the Ter site and was almost linear; ii) clear (14 cMAGs), whereby the curve displayed clear symmetry but also a few minor non-Ter site peaks (few thousand bp); iii) decent (53 cMAGs), whereby the most significant peak of the curve was the Ter site, but several minor peaks also existed; iv) poor (10 cMAGs), whereby the curve exhibited distinct ascending and descending regions but no symmetry or had significant peaks other than the Ter site; and v) no-pattern (8 cMAGs), whereby the curve presented no GC-skew pattern for Ter or Ori sites. The GC-skew class annotation is provided in **Supplementary Table S4**, and GC-skew plots are presented in **Extended Data Fig. 4**.

### SNV analysis

We aligned cMAGs to HRGM species representatives and found conspecific HRGM species with ANI > 0.95 and maximum alignment coverage > 0.6. For cMAGs with multiple conspecific HRGM species, we selected only the species with the highest similarity index (ANI × maximum alignment coverage). To obtain a reliable SNV density percentile value of cMAG, we performed subsequent SNV analysis only if there were > 100 conspecific genomes for the HRGM species cluster. We aligned cMAG and HRGM genomes (non-representative) against representative HRGM species using nucmer, and filtered the best bidirectional alignments using a delta-filter with *-r* and *-q* options. Next, to eliminate indels, SNVs were identified using *show-snps* in the *mummer* package with *-I* options. The overall procedure is depicted in **Extended Data Fig. 5**.

### Conspecific isolated or complete genomes for cMAGs

We compiled isolated genome sequences from HRGM^3^, hGMB^31^, and the recently updated NCBI genome databases^16,32^. Then, we aligned cMAGs to isolated genomes with nucmer and delta-filter, as described in the redundancy removal step. We defined cMAGs with conspecific isolated genomes as having at least one isolated genome, ANI > 0.95 and maximum alignment coverage > 0.6. For such cMAGs, we manually searched whether the conspecific isolate was complete in the NCBI genome database.

### Reconstruction of the phylogenetic tree

We predicted bac120 or arc122 proteins using the GTDB-Tk identify module^25^ and performed multiple sequence alignment with the GTDB-Tk align module. The tree was calculated using IQ-Tree v2.1.3^33^ and visualized with ITOL v6^34^.

### Examining the taxonomic distinctiveness of RF39 and TANB77 orders

To investigate the phylogenetic distinctiveness of RF39 and TANB77 orders, we gathered HRGM species and cMAGs within the Bacilli and Clostridia classes, which were their respective parental clades. We then constructed a phylogenetic tree of Bacilli and Clostridia, and selected the top 100 closest genomes of RF39 and TANB77 orders based on the average maximum likelihood distance. For RF39 and TANB77 orders and each of their close neighboring genomes, we predicted their proteins with Prokka^35^ and annotated protein functions with eggNOG-mapper v2.1.6^36^. Hierarchical clustering was performed using KEGG Orthology and KEGG Pathway^37^ profiles based on Euclidean distance.

### Genome coverage by the short-read assembled genome catalog

We measured the coverage of cMAGs to identify hard-to-assemble genome regions by short-read assembly. First, we aligned conspecific genomes to each cMAG using nucmer, and identified the best alignments with delta-filter using the *-r* parameter. We then calculated the genome coverage for every 1-kbp bin, *B*, of cMAG as follows:

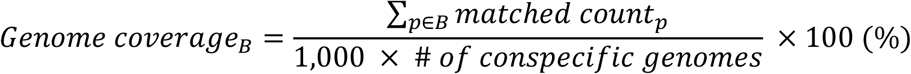

where *p* indicates the single nucleotide position in *B*.

To investigate the characteristics of low-coverage regions, we compared the genome coverage of genome bins containing rRNAs or genomic islands with other regions. We predicted the 5S, 16S, and 23S rRNA regions using barrnap (as described in the first filtration section). In addition, we annotated cMAGs with Prokka v1.14.6^35^ and identified genome island regions using IslandViewer 4^38^. Proteins located within genome islands were predicted by prodigal v2.6.3^39^, and we annotated their eggNOG ortholog and function using eggNOG-mapper 2^36^.

## Data availability

HiFi metagenomic sequencing data generated in this study were deposited in the Sequence Read Archive under codes PRJNA798244. The entire 102 cMAGs sequences and their GC-skew and genome coverage plots are available at https://doi.org/10.5281/zenodo.5996768

## Author contributions

C.Y.K. and I.L. conceived and designed the study. C.Y.K. and J.M. performed the bioinformatics analysis on metagenomic data and formulated the study hypothesis. I.L. supervised the bioinformatics analysis. All authors contributed to the writing of the manuscript.

## Acknowledgements

This research was supported by the National Research Foundation funded by the Ministry of Science and ICT (2018R1A5A2025079 and 2019M3A9B6065192) and supported in part by the Brain Korea 21 (BK21) FOUR Program. HiFi sequencing was provided by the SMRT Grant of MdxK, Macrogen, and PacBio.

## Extended Data Figures

**Extended Data Fig.1.**
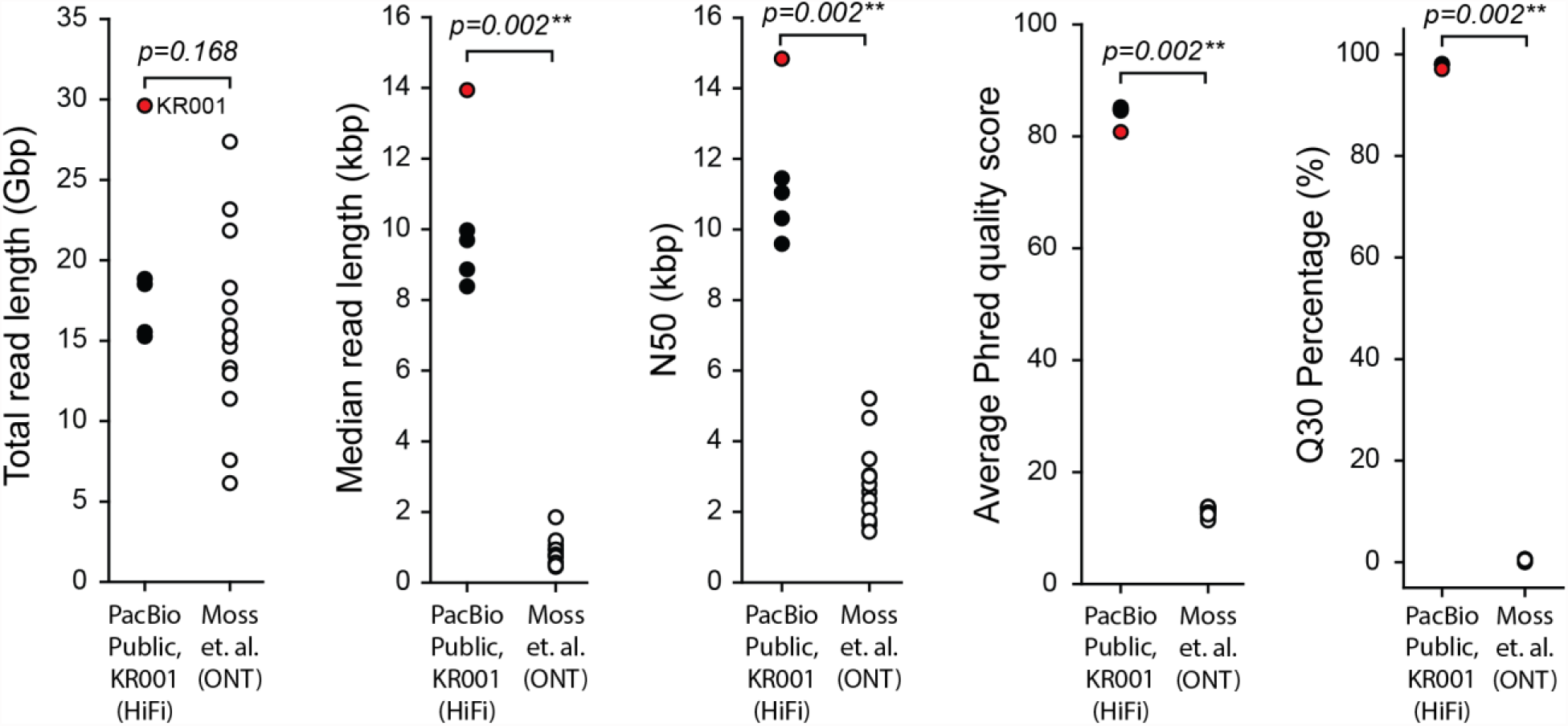
Comparison of high-accuracy long-read (HiFi) and Oxford nanopore technology (ONT) sequencing data on human feces. Total read length, median read length, N50, average Phred33 score, and Q30 percentage of HiFi and ONT data were compared using a two-sided Mann–Whitney U test. The red data point represents the in-house HiFi sequencing data from a healthy Korean donor (\*\**P* < 0.01).

**Extended Data Fig.2.**
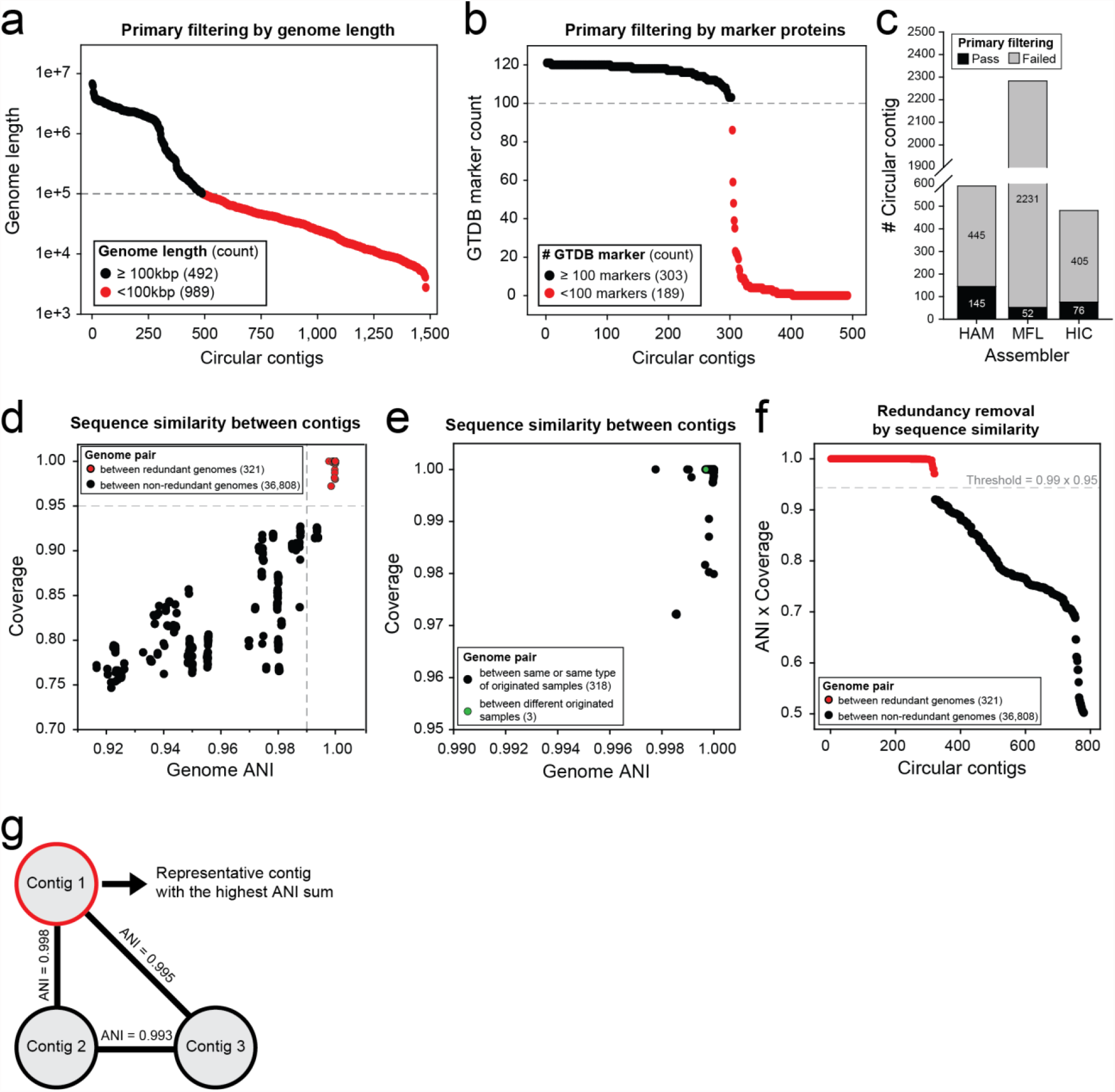
Filtering of circular contigs by biological priors and removal of redundant contigs. **a, b**, Filtering of circular contigs by length (a) and number of genome taxonomy database (GTDB) marker proteins (b). Contigs that passed the criteria are denoted in black; whereas those that failed to meet the criteria are denoted in red. **c**, Number of circular contigs before and after the first filtration step for each assembler. **d**, Aaverage nucleotide identity (ANI) and maximum alignment coverage between pairwise circular contigs. Only contig pairs with ANI > 0.9 and maximum alignment coverage > 0.7 are shown in the plot. Contig pairs between redundant contigs with ANI > 0.99 and coverage > 0.95 are highlighted in red. **e**, Highly similar contig pairs with ANI > 0.99 and maximum alignment coverage > 0.95. Most contig pairs were between contigs from the same or the same type of sample (black). **f**, Genome similarity index (ANI × maximum alignment coverage) for contig pairs. The horizontal dashed line represents the score threshold for redundancy removal. Red data points represent contig pairs subjected to redundancy removal. **g**, Conceptual diagram showing how the representative contig was selected from the redundant contig group.

**Extended Data Fig.3.**
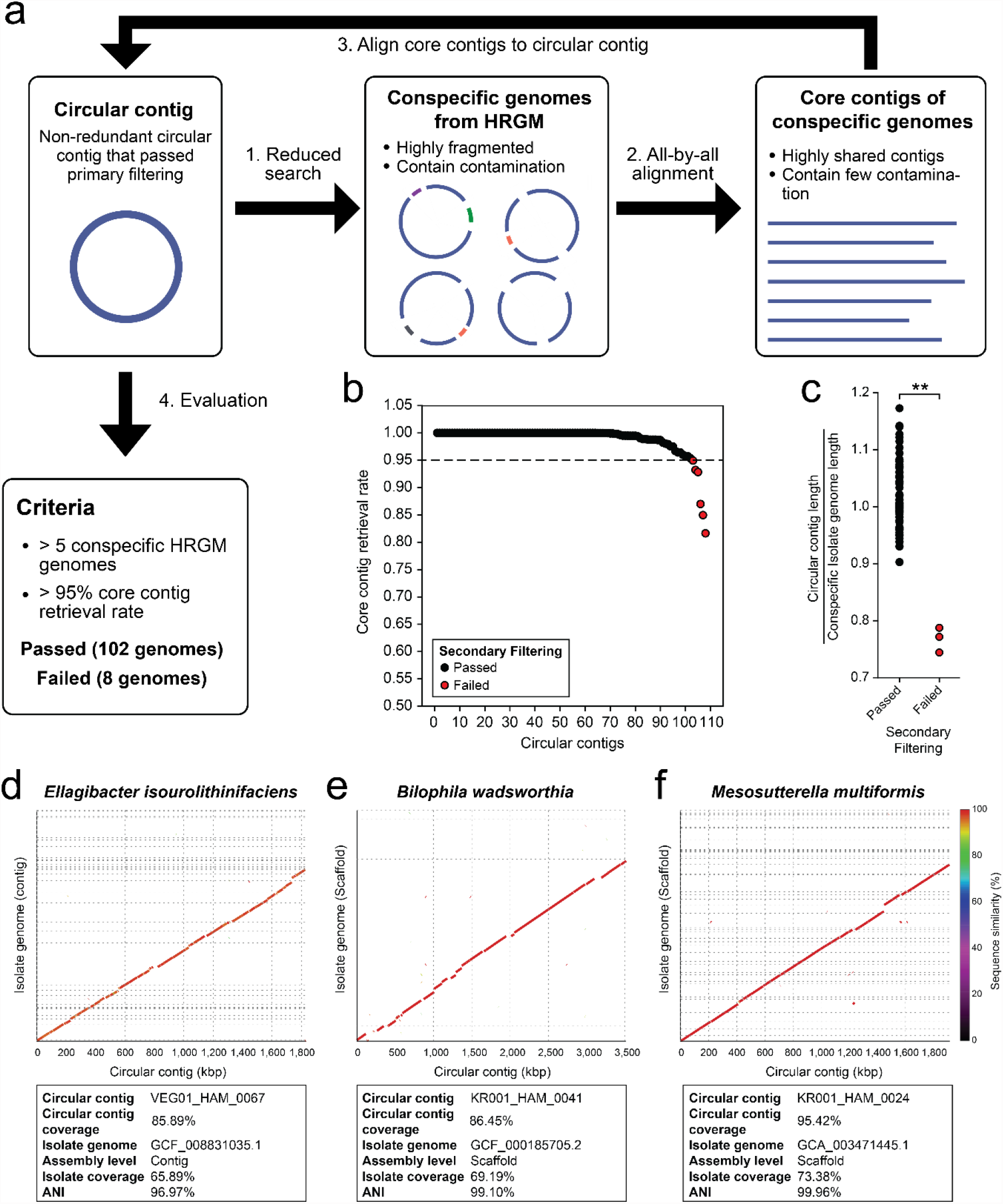
Filtering circular contigs by congruency with conspecific genomes. **a**, Schematic diagram illustrating the filtering pipeline. **b**, Filtered circular contigs by core contig retrieval rate. Contigs that passed the filtering threshold are denoted in black; whereas those that failed to pass the threshold are denoted in red. **c**, Relative genome length of circular contigs compared to their conspecific isolated genomes. Relative genome lengths of contigs that passed or did not pass the criteria were compared using the two-sided Mann–Whitney U test. **d–f**, Genome alignment plot between circular contigs that did not pass the cirteria and their closest isolated genomes. Genome alignment plot for *Ellagibacter isourolithinifaciens* (d), *Bilophila wadsworthia* (e), and *Mesosutterella multiformis* (f).

**Extended Data Fig.4.**
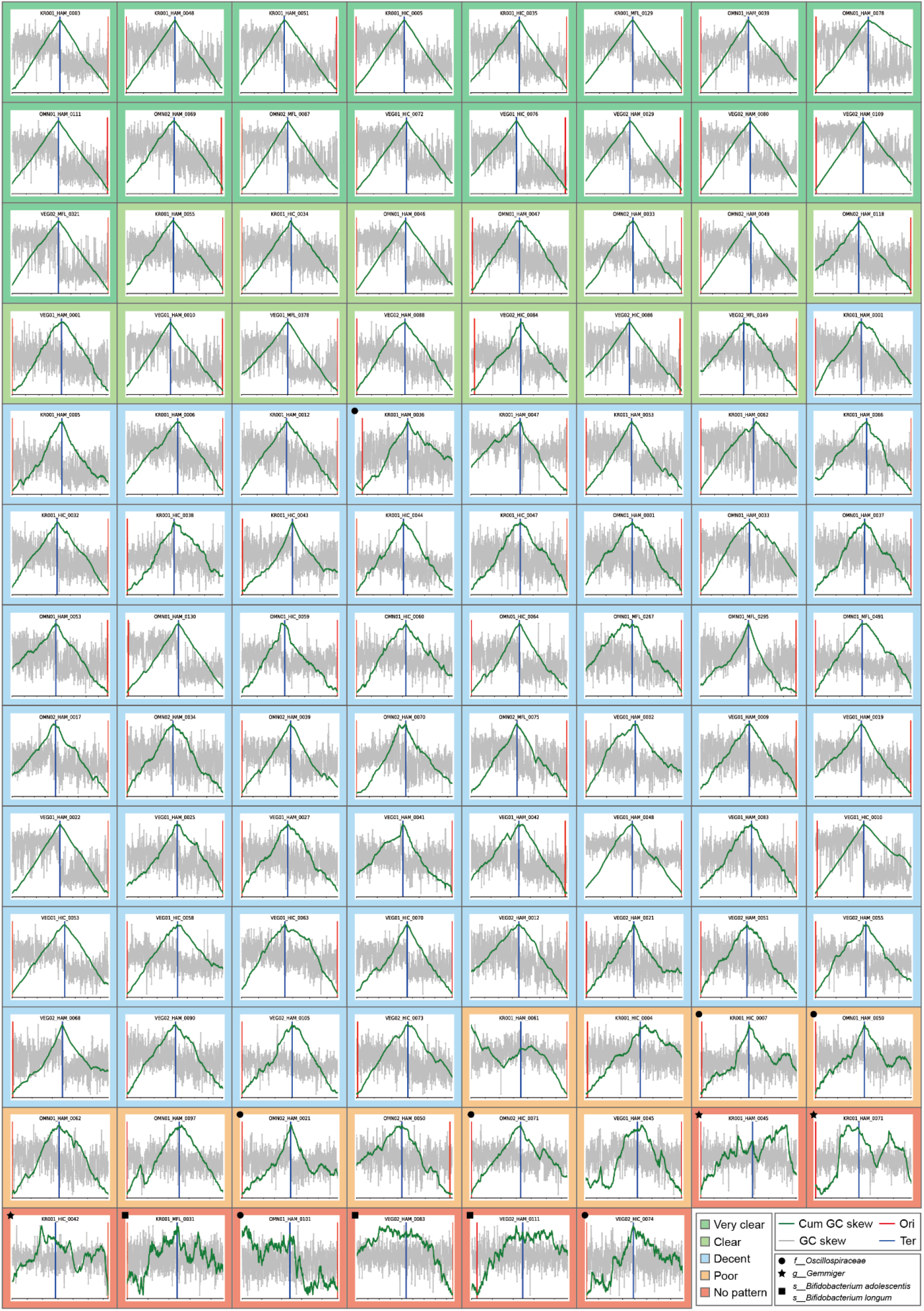
Five classes of circular contigs based on cumulative GC-skew curves. Different background colors represent different classes. Bacterial lineages repeatedly observed in the poor or no-pattern classes are marked.

**Extended Data Fig.5.**
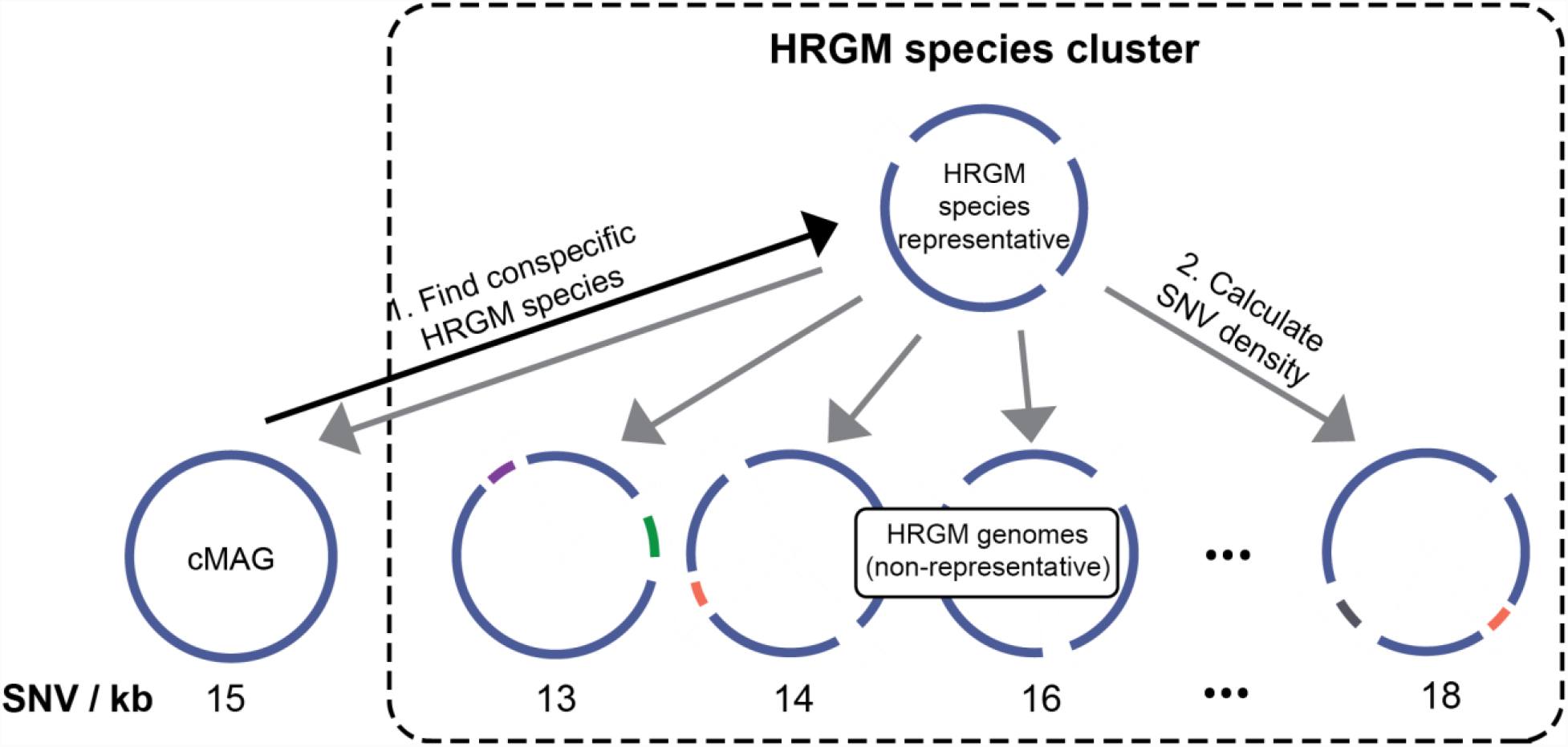
Schematic diagram representing the pipeline for comparing single nucleotide variation (SNV) density of complete metagenome-assembled genomes (cMAGs) to their conspecific genomes.

**Extended Data Fig.6.**
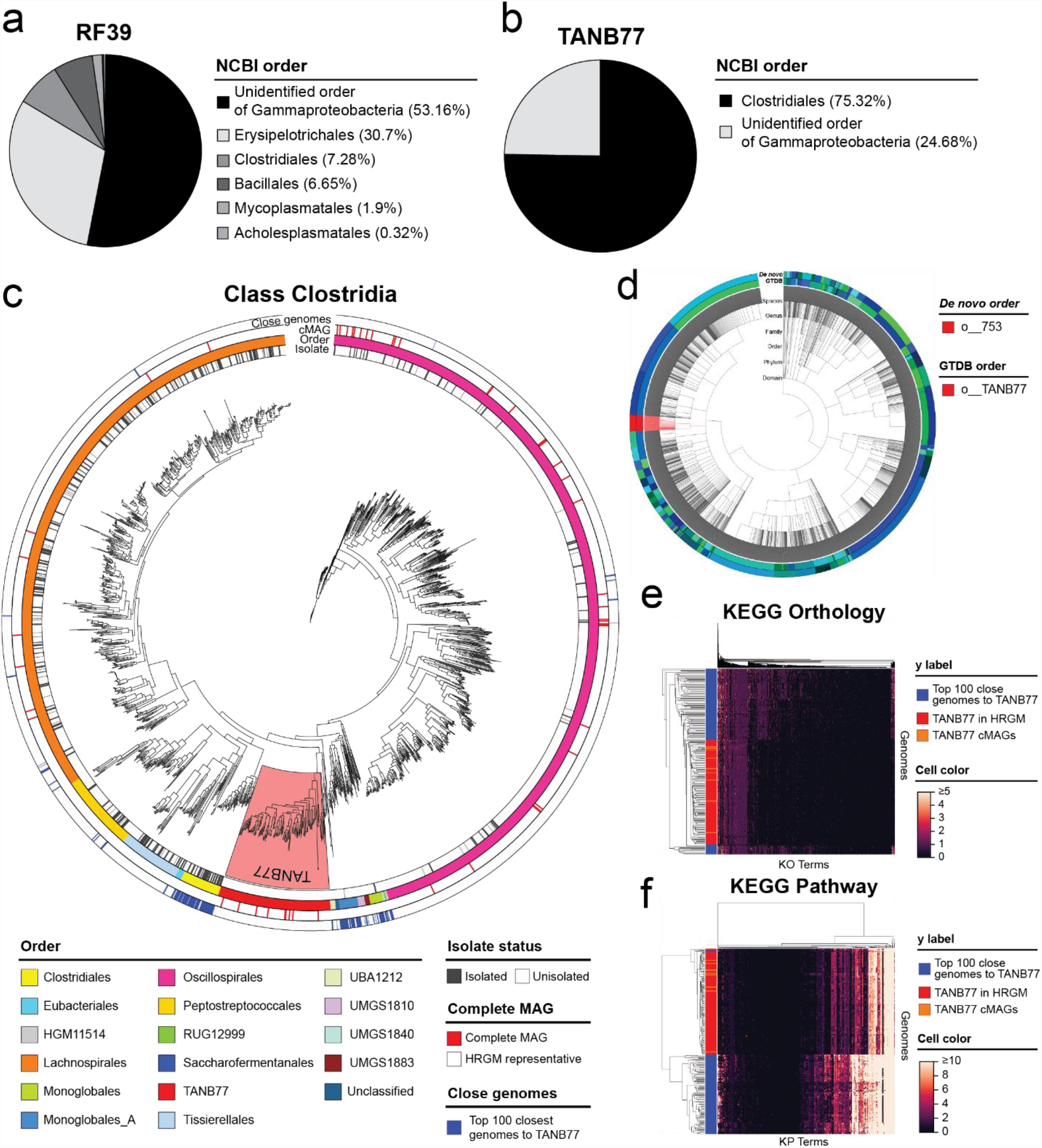
Complete genomes of entirely unculturable bacterial orders. **a, b**, Proportion of NCBI orders defining RF39 (a) and TANB77 (b). **c**, Maximum likelihood phylogenetic tree of human reference gut microbiome (HRGM) species and HiFi cMAGs in the Clostridia class. The red-highlighted area represents the TANB77 order. Isolated status, order, HiFi cMAG, and top 100 closest genomes to the TANB77 order are annotated from the innermost to the outermost circle, respectively. **d**, Phylogenetic tree of 4,545 bacterial species of HGM. The inner circle annotates the order according to the GTDB; whereas the outer circle represents the order according to HGM *de novo* classification. TANB77 and o_047 orders are highlighted in red. **e, f**, Hierarchical clustering of heatmaps representing KEGG Orthology (e) and KEGG Pathway (f) profiles of 129 TANB77 (red and orange rows) and the top 100 closest genomes to TANB77 (blue rows).

